# Dissolved oxygen concentrations influence microbial diversity, abundance and dominant players in an oxygen minimum zone

**DOI:** 10.1101/2025.10.12.681960

**Authors:** Kaitlin R. Dombroski, Lauren Gillies Campbell, Robert M. Morris, Olivia U. Mason

## Abstract

Expansion of marine global oxygen minimum zones (OMZs) can have profound impacts on resident macrofauna. Less obvious is the influence OMZs will have on the diversity and abundance of planktonic microbes. This is particularly true in understudied OMZs such as the northern Benguela Upwelling System (nBUS). Here, we analyzed the influence of oxygen concentrations on the microbial community in the nBUS OMZ using 16S rRNA gene (iTag) sequence data. In the nBUS oxygen was a primary driver influencing microbial community structure and diversity. Diversity was highest in dysoxic samples and lowest in suboxic samples, which was primarily due to changes in community evenness in relationship to oxygen concentrations. For example, evenness decreased in suboxic samples due to oscillations in the abundance of microbial groups such as *Thioglobaceae* (SUP05), which was found to be the most abundant microbe in the nBUS OMZ and significantly increased in abundance as oxygen decreased. This finding prompted an analysis of 217 publicly available medium to high quality *Thioglobaceae* genomes, including cultured representatives, from the nBUS and other OMZs. Genome annotation of these *Thioglobaceae* indicated important roles in carbon cycling, sulfur oxidation and denitrification. Importantly, few *Thioglobaceae* possess the genetic potential to carry out complete denitrification, as most lack the gene that codes for nitrous oxide reductase (NosZ), which converts nitrous oxide (N₂O), a potent greenhouse gas, to nitrogen gas. As OMZs expand in size and severity, decreasing microbial diversity and a concomitant increase in *Thioglobaceae* abundances, could lead to enhanced N₂O production through incomplete denitrification.

**Importance:** Here, we found that microbial diversity decreased significantly with declining oxygen concentrations in the nBUS OMZ. However, one microbial group, the *Thioglobaceae*, was most abundant when oxygen was lowest. This group is able to support its growth through sulfur oxidation using either oxygen or nitrate (denitrification). A comprehensive genomic analysis of *Thioglobaceae* in the nBUS and in the global ocean revealed that few have the capacity to carry out complete denitrification, with the final step in this process often missing in these genomes. This incomplete pathway is consequential, as it can be an important source of nitrous oxide, particularly in marine OMZs such as the understudied nBUS. Collectively, this study provides new information on an OMZ, and definitively links an important microbial pathway, incomplete denitrification, to a particular and highly abundant group of microbes that appear to have a strong response to declining oxygen concentrations in the marine environment.

## Introduction

Eastern Boundary Upwelling Ecosystems (EBUEs) are some of the most productive regions of the ocean, contributing up to 11% of global primary production (Chavez & Toggweiler, 1995; Arévalo-Martínez et al., 2019). Among EBUEs, the Benguela Upwelling System (BUS) has been reported to be the most productive, accounting for up to 40% of global EBUE production (Carr, 2002; Monteiro, 2010). High productivity in the region is attributed to year-round upwelling (Carr, 2002; Arévalo-Martínez et al., 2019) facilitated by trade winds that displace surface waters and transport cold, nutrient-rich water to the surface (Martin et al., 2017). Specifically, the northern Benguela Upwelling System (nBUS) lies off the coast of Namibia, between the Angola-Benguela front and the Lüderitz upwelling cell (Mohrholz et al., 2008; Martin et al., 2017). The nBUS ecosystem is further influenced by the South Atlantic Central Water (SACW) mass, which brings oxygen-depleted and nutrient-rich water south from Angola (Angola Current), and the nutrient-poor, oxic Eastern South Atlantic Central Water (ESACW) mass, which flows north from South Africa (Agulhas Current). Thus, large scale and regional circulation patterns determine the composition of the upper central water layer and subsequently the oxygen balance over the Namibian shelf (Mohrholz et al., 2008).

In EBUEs, and more broadly in marine oxygen minimum zones (OMZs), high productivity leads to enhanced aerobic respiration and oxygen depletion at depth (Arévalo-Martínez et al., 2019). OMZs are defined as regions where dissolved oxygen (DO) is less than 62 μM/kg or 2 ml/L (Ulloa et al., 2012; Wright et al., 2012; Roux et al., 2014; Canfield & Kraft, 2022). OMZs are expanding due to rising ocean temperatures, which decreases oxygen solubility and reduces water column ventilation (Keeling et al., 2010; Rabalais et al., 2010; Helm et al., 2011; Wright et al., 2012). Declining oxygen concentrations shape community structure and can lead to shifts in both macro and microfauna (Breitburg et al., 2018). Changes in microbial community composition associated with low DO levels, especially under dysoxic (DO ranging from 0.2 to 2.0 ml/L) conditions, have been documented (Aldunate et al., 2018; Spietz et al., 2015). For example, bacterial diversity and evenness have been reported to decrease within hypoxic zones, particularly in OMZs (Beman & Carolan, 2013; Bryant et al., 2012). In contrast, bacterial richness has an inverse relationship with DO (Beman & Carolan, 2013; Spietz et al., 2015). Specifically, Beman and Carolan (2013) reported that richness increased with declining DO and decreasing depth. Changes in DO have the potential to influence microbial diversity, specifically richness, which could alter the function of the microbial community and the prevalence of key metabolic pathways (Fernandes et al., 2020).

Taxonomic surveys have identified consistent trends in microbial community composition along DO gradients, from the anoxic core to the boundaries of OMZs (Beman and Carolan, 2013). In oxygen-depleted waters, an increase in microaerophilic and anaerobic metabolisms, such as denitrification, anaerobic ammonium oxidation, and sulfate reduction has previously been noted (Long et al., 2021). *Thermoproteota*, *Pseudomonadota*, *Nitrospinota*, SAR324, and *Nanobdellota* are among the dominant taxa in OMZs (Zaikova et al., 2010; Beman and Carolan, 2013; Bandekar et al., 2018; Beman et al., 2020; Pajares et al., 2020; Vuillemin et al., 2022; Anstett et al., 2023). Although ammonia oxidizing archaea, in the *Thermoproteota*, are frequently identified in low DO environments (Lam et al., 2007; Beman et al., 2008; Molina et al., 2010; Belmar et al., 2011; Tolar et al., 2013; Gillies et al., 2015; Campbell et al., 2019), OMZs are typically dominated by *Pseudomonadota* (Wright et al., 2012), particularly sulfur-oxidizing members of the family *Thioglobaceae* (Lavik et al., 2009; Walsh et al., 2009; Glaubitz et al., 2013), which has historically been referred to as SUP05, but is now classified as a genus in the *Thioglobaceae*. For example, *Thioglobaceae* abundances were sufficiently high that detoxification of sulfidic waters in the nBUS was attributed, in part, to this microbial group (Lavik et al., 2009).

Chemoautotrophic *Thioglobaceae* use energy gained through aerobic or anaerobic oxidation of reduced sulfur to fix inorganic carbon. In the absence of oxygen, they use nitrate as a terminal electron acceptor (Shah et al., 2019). A recent study found that *Candidatus Thioglobus autotrophicus* strain EF1 simultaneously respired oxygen and nitrate when DO concentrations were low (<5 μM) and while nitrate was available (Mattes et al., 2021). This led to higher growth rates, enhanced carbon fixation, and elevated gene expression (Mattes et al., 2021). Importantly, previous experiments with *Ca. T. autotrophicus* revealed that nitrous oxide (N₂O) was produced as a result of incomplete denitrification (Shah et al., 2017). Strain EF1 does not code for nitrous oxide reductase (NosZ), the enzyme required to complete the final step of denitrification, in which N₂O is reduced to nitrogen gas (Shah et al., 2017). This is consistent with previous reports of an incomplete denitrification pathway in *Thioglobus*, a genus within the *Thioglobaceae* (Walsh et al., 2009; Schunck et al., 2013). OMZs account for approximately 60% of marine N₂O emissions (Suntharalingam et al., 2000; Ji et al., 2018; Yang et al., 2020), and the high abundance of *Thioglobaceae* in OMZs (Walsh et al., 2009), coupled with laboratory experiments showing that *Ca*. *T. autotrophicus* produces N₂O (Shah et al., 2017), suggests that chemoautotrophic members of the *Thioglobaceae* may be an important source of N₂O in OMZs.

As the nBUS OMZ is relatively understudied compared to other EBUEs, we used 16S rRNA gene sequencing (iTag) to analyze shifts in microbial diversity across oxygen gradients. The iTag dataset revealed a dominance of *Thioglobaceae*; therefore, a set of *Thioglobaceae* genomes was compiled and analyzed to determine the metabolic function of this group in the nBUS and, more broadly, in the global ocean. Specifically, a comprehensive analysis of all publicly available *Thioglobaceae* genomes, which included cultured representatives with complete genomes and genomes from metagenome assemblies and single amplified genomes that were assembled to a minimum of 50% complete with less than 10% contamination (217 genomes total; 78% average genome completeness) was carried out. These genomes represent microbes sampled from the nBUS and the global ocean and were evaluated to resolve the metabolic function of *Thioglobaceae* in key biogeochemical processes, including their potential to produce the potent greenhouse gas N₂O within OMZs.

## Results

### Microbial Community Diversity

Analysis of the ASV data from the nBUS (Fig. 1A) revealed that microbial community diversity (Shannon diversity) was highest in dysoxic samples and lowest in suboxic samples (Fig. 1B). Further, diversity was significantly lower in the oxic and suboxic samples compared to the dysoxic samples (Fig. 1B). Similarly, species richness (Chao1) was highest in dysoxic samples and significantly lower in both oxic and suboxic samples. (Fig. 1C). Community evenness (Pielou e) was highest in the oxic and dysoxic samples and significantly lower in the suboxic samples (Fig. 1D).

**Figure 1.**
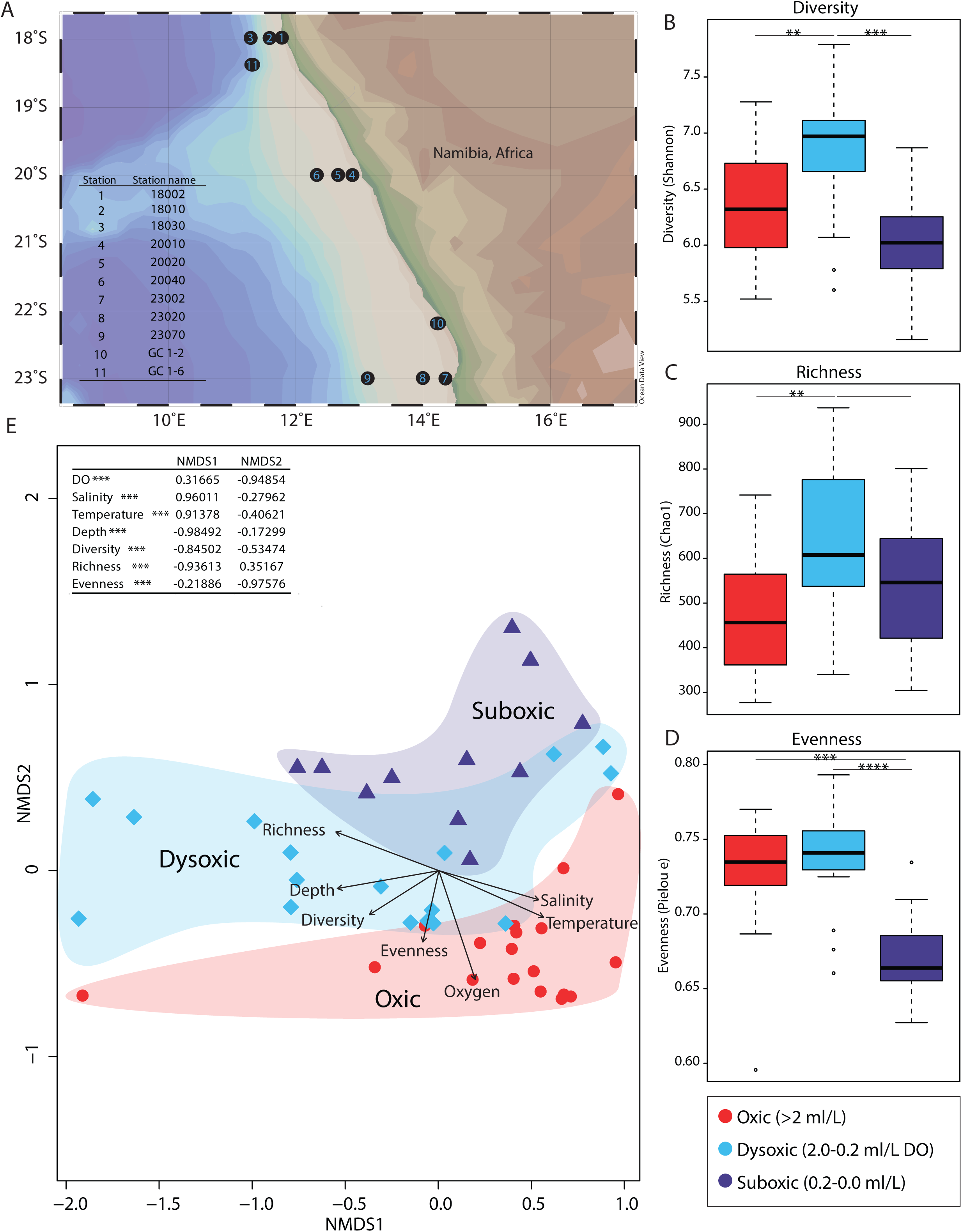
(A) Sampling stations off the coast of Namibia in the northern Benguela upwelling system (nBUS). (B, C, D) Boxplots show diversity metrics of 16S rRNA gene sequence data. Microbial diversity, richness, and evenness were compared across oxic, dysoxic and suboxic conditions. (E) Non-metric multi-dimensional scaling (NMDS) ordination of 16S rRNA gene sequence data. Environmental data was overlain on the ordination, with variable correlations with ordination axes shown in the table in the figure. Corrected p-values were symbolized by ** ≤ 0.01, *** ≤ 0.001, and **** ≤ 0.0001 in the figures.

### Microbial Community Structure

To identify the influence of oxygen concentrations on community structure, including changes in diversity, richness, and evenness, normalized ASV data was evaluated using NMDS ordination (Fig. 1E). Environmental data and diversity metrics were overlain onto the ordination, revealing that dissolved oxygen (DO) and depth were the most significant drivers structuring microbial communities, with samples from similar DO concentrations and depth clustering together (Fig. 1E). Sample clustering based on DO concentrations were significantly different, as determined using PERMANOVA. In addition to depth and DO, other environmental factors such as temperature and salinity were also significantly correlated with NMDS axes (Fig. 1E).

Diversity metrics were also overlain on the ordination (Fig. 1E), the results of which agreed with the variation in diversity, richness, and evenness discussed previously (Figs. 1B-D). For example, diversity, richness, and evenness were highest in dysoxic samples, while diversity and evenness were lowest in suboxic samples and richness was lowest in oxic samples (Figs. 1B-E).

Of the 5,463 ASVs in all samples, SIMPER analysis indicated that 138 contributed significantly to the observed DO clusters in the NMDS ordination. These ASVs had significantly different abundances based on DO, as determined by a Kruskal-Wallis test. These ASVs were predominantly *Thioglobaceae* in the *Gammaproteobacteria* (Fig. 2), *Poseidoniales*, and *Nitrosopumilaceae*. In particular, an ASV classified as *Thioglobaceae* was the most significant microbe contributing to the observed clustering when comparing oxic to suboxic samples and oxic to dysoxic clusters. Further, the relative abundance of members of the *Thioglobaceae* comprised a minimum of 6% to a maximum of 40% of the entire microbial community in samples (Fig. 2B) and was observed in all samples; thus, *Thioglobaceae* were the most abundant microbial group in the nBUS.

**Figure 2.**
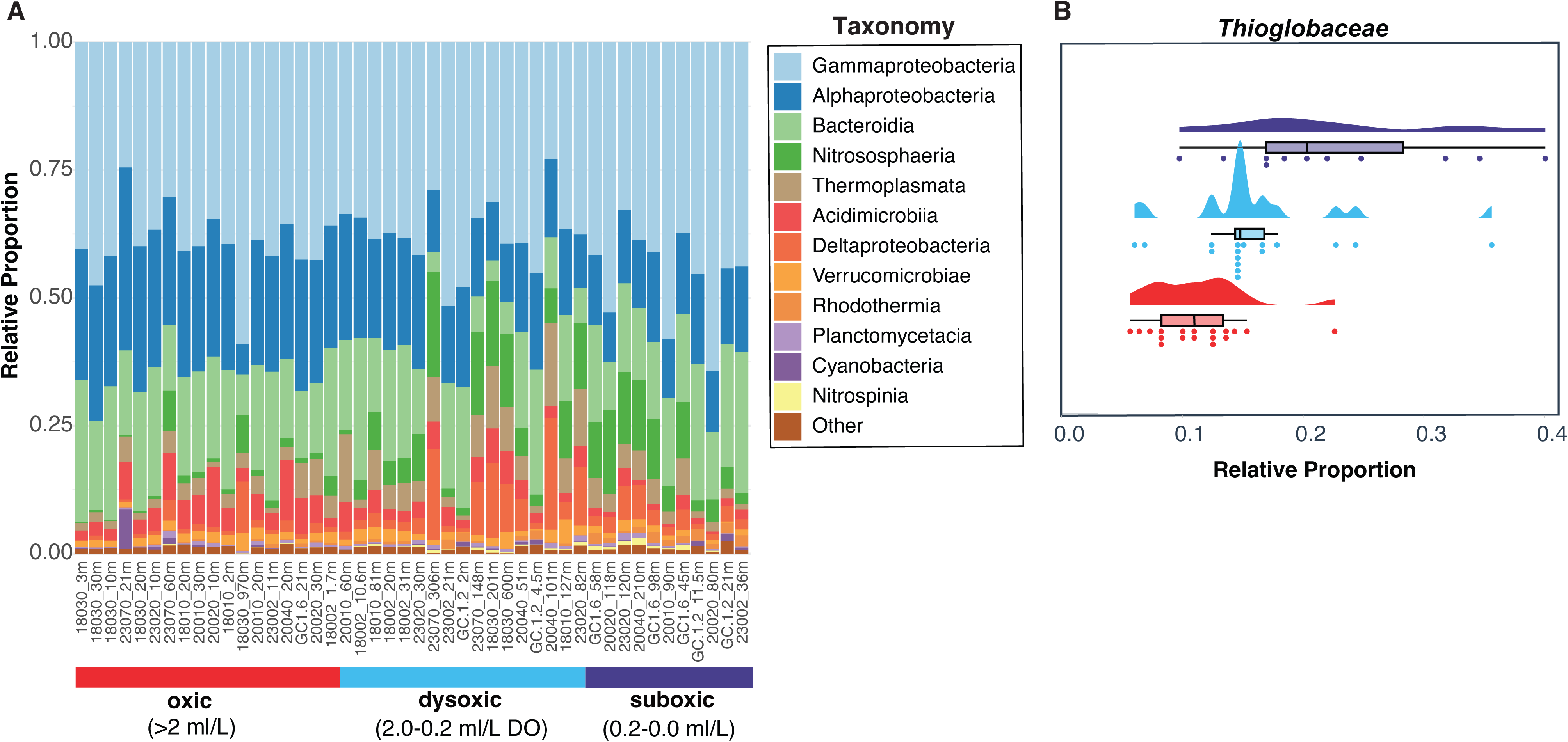
(A) Bar graph of 16S rRNA gene data showing relative proportions of phyla, and in the case of *Pseudomonadota*, classes. Only the most abundant groups are shown, with less abundant phyla grouped as “Other”. Samples are sorted by DO concentration (suboxic, dysoxic, and oxic). (B) The raincloud plot shows the relative abundance of *Thioglobaceae,* from 16S rRNA gene sequence data in relationship to oxygen concentrations.

At higher taxonomic levels, the *Thioglobaceae* (*Pseudomonadota*), *Alphaproteobacteria* (*Pseudomonadota*), *Bacteroidota, Poseidoniales* (*Thermoplasmatota*), and *Nitrosopumilus* (*Thermoproteota*) dominated in the nBUS (Fig. 2). Some of these groups were nearly invariant across the DO gradient, such as *Poseidoniales*, while other groups were highly variable. For example, *Thioglobaceae* decreased in abundance with increasing DO concentrations (Fig. 2), as did *Nitrosopumilaceae*.

### Genome Classification, Quality, and Average Nucleotide Identity

As the most abundant group in the iTag data, *Thioglobaceae* genomes were subsequently compiled and analyzed to evaluate the functional roles of this group in the environment. The 217 genomes were medium to high quality and represented cultured and uncultured microbes in the genera *Thioglobus*, SUP05, *Pseudothioglobus*, SZUA-1055, *Thiodubiliella*, DUCF01, and JAAOIF01 (Supp. Fig. 1). To link *Thioglobaceae* in the iTag data to genomes, BLAST analysis was used to compare 16S rRNA genes to determine sequence similarity between *Thioglobaceae* ASVs and genomes. Of the 96 genomes that coded for 16S rRNA genes, 93 had a BLASTn hit from the iTag data, and 91 were 100% similar to the *Thioglobaceae* ASVs. When the iTag data was converted to relative abundances, these *Thioglobaceae* ASVs represented a minimum of 89% and a maximum of 100% (average 98%) of all identified *Thioglobaceae* in the iTag dataset.

The 217 genomes discussed herein were sequenced from microbes sampled in the global ocean, including samples from the nBUS (NCBI accession PRJNA930832). These genomes also include the first cultured *Thioglobaceae* (referred to as SUP05) representative, *Ca. T. autotrophicus* strain EF1 (Shah & Morris, 2015), and the recent isolates EF2 and EF3 (Morris & Mino, 2024) from the Effingham Inlet, British Columbia, as well as the genome of *Ca*.

*Thioglobus,* strain NP1, from the North Pacific subpolar gyre (Spietz et al., 2019). Additional genomes were from cultured and uncultured microbes sampled in the Peru upwelling region OMZ, such as *Ca. T. perditus* (Callbeck et al., 2018), and numerous *Thioglobaceae* (SUP05) sampled from seawater off the Chilean coast OMZ and in the seasonally anoxic Saanich Inlet in British Columbia (Walsh et al., 2009).

The 217 genomes analyzed here had an average genome completeness of 78%, with an average of 1.6% contamination. The 16S rRNA gene was encoded in 96 of these genomes.

Average nucleotide identity (ANI) analysis of these genomes revealed ANI values ranging from > 75% to 99% in similarity within genera and in some instances across the *Thioglobus*, SUP05, SZUA-1055, and *Pseudothioglobus* genera (Supp. Fig. 1). Generally, ANI values were <75% when comparing genomes across the different taxonomic groups (Supp. Fig. 1).

### Carbon fixation and utilization

Analysis of carbon fixation pathways revealed that 13% (29 of 217) of *Thioglobaceae* genomes encoded a complete carbon fixation pathway via the canonical Calvin-Benson cycle (CB; Fig. 3), as defined in Hudson et al. (2024). SUP05 and *Thioglobus* genera were identified as the only clades whose members completed carbon fixation via the canonical CB cycle, as they encoded all genes in this pathway. This included the bifunctional enzyme fructose-1,6- bisphosphatase/sedoheptulose-1,7-bisphosphatase (F/SBpase; Fig. 3), which hydrolyzes FBP and SBP. Of the 217 genomes, 60 in the SUP05, *Thioglobus*, *Pseudothioglobus*, SZUA-1055, *Thiodubiliella*, DUCF01, and JAAOIF01 genera specifically lacked only F/SBpase, while still encoding the remainder of the CB cycle. It is notable that the EF1 strain, as well as the EF2 and EF3 isolates (Shah & Morris, 2015; Morris & Mino, 2024), all lacked only F/SBpase. The remaining 128 genomes, including a mix of all but JAAOIF01 and DUCF01, encoded steps within the CB cycle but lacked a complete pathway.

**Figure 3.**
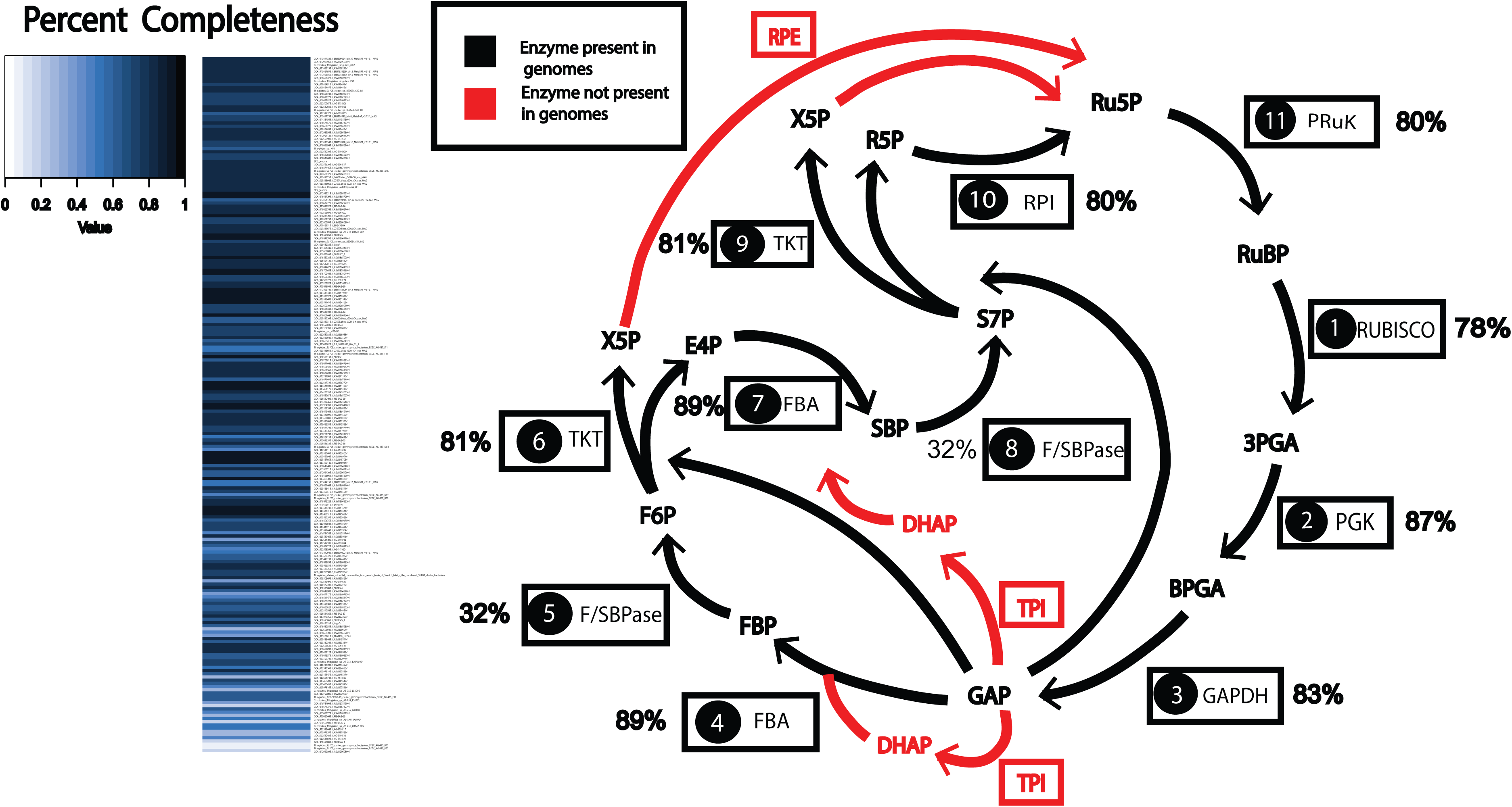
Carbon fixation pathway via the canonical Calvin-Benson cycle. The 11 key enzymes in the canonical Calvin-Benson cycle are in order as follows: 1) ribulose-bisphosphate carboxylase (RuBisCO) 2) phosphoglycerate kinase (PGK) 3) glyceraldehyde-3-phosphate dehydrogenase (GAPDH) 4) fructose-bisphosphate aldolase (FBA) 5) fructose-1, 6- bisphosphatase/sedoheptulose-1, 7-bisphosphatase (F/SBpase) 6) transketolase (TKT) 7) fructose-1,6-bisphosphatase (FBA) 8) fructose-1, 6-bisphosphatase/sedoheptulose-1, 7- bisphosphatase (F/SBpase) 9) transketolase (TKT) 10) ribose 5-phosphate isomerase (RPI) 11) phosphoribulokinase (Ru5P). The percentages shown next to the enzyme represent the percent of genomes coding for each enzyme. Enzymes shown in black were present in the genomes, while enzymes shown in red indicate absence of those enzymes in the genome. The products of each step of the pathway are shown in black in between the arrows. The percent completeness of this pathway in each genome is shown on the heatmap to the left of the cycle diagram.

Carbon utilization was also analyzed in all 217 genomes, revealing that the tricarboxylic acid cycle (TCA) was encoded in all genomes but with varying degrees of pathway completion. For example, key components of the TCA cycle, such as 2-oxoglutarate dehydrogenase (K00164), were lacking in 70% (150 of 217) of the genomes, including most *Thioglobus* and all SUP05. This suggests that the majority of *Thioglobaceae*, *Thioglobus* and SUP05 in particular, carry out a modified TCA cycle. In contrast, nearly all *Pseudothioglobus* coded for this enzyme. In 62 genomes that lacked 2-oxoglutarate dehydrogenase genes, malate dehydrogenase (K00024), another key component of the TCA cycle, was also missing. Genomes lacking both genes were predominantly classified as *Thioglobus*, however, all other *Thioglobaceae* genera were also represented in this group.

### Aerobic and Anaerobic Respiration

#### Aerobic Respiration

Over 70% (152 of 217) of the genomes encoded cytochrome c oxidase subunits I-III (CoxABC), a low oxygen affinity enzyme. Nearly 60% (127 of 217) of the genomes analyzed encoded cbb3-type cytochrome c oxidase subunits I-II, a high oxygen affinity enzyme. All *Thioglobaceae* clades had representatives encoding this enzyme. Both low and high affinity enzymes were encoded in 98 genomes. In total only 15% (33 of 217) of the genomes lacked either low or high affinity enzymes.

#### Anaerobic Respiration (Denitrification)

In addition to aerobic respiration, 37% (80 of 217) of the genomes, from both cultured and uncultured representatives, encoded the necessary genes to carry out denitrification, with varying degrees of pathway completion. This included *Thioglobus*, *Pseudothioglobus*, and SUP05, with fewer from DUCF01, *Thiodubiliella*, and SZUA-1055. Of the 80 genomes, only six *Thioglobus*, including *Ca. T. perditus* (Callbeck et al., 2018), encoded all the necessary genes for complete denitrification, including nitrate reductase (*napAB*), respiratory nitrate reductase *(narGHI*), nitrite reductase subunit K (*nirK*) (only one genome encoded *nirS*), nitric oxide reductase (*norBC)*, and nitrous oxide reductase (*nosZ)* (Fig. 4). The final step of denitrification, which is the conversion of nitrous oxide (N₂O) to nitrogen gas (N₂), is mediated by *nosZ* (Fig. 4), which was lacking in 14 genomes that encoded all other enzymes in the denitrification pathway (Fig. 4). These genomes included 13 classified as SUP05 and one classified as *Thioglobus.* The remaining 60 genomes encoded some steps of the denitrification pathway but not for complete denitrification (Fig. 4).

**Figure 4.**
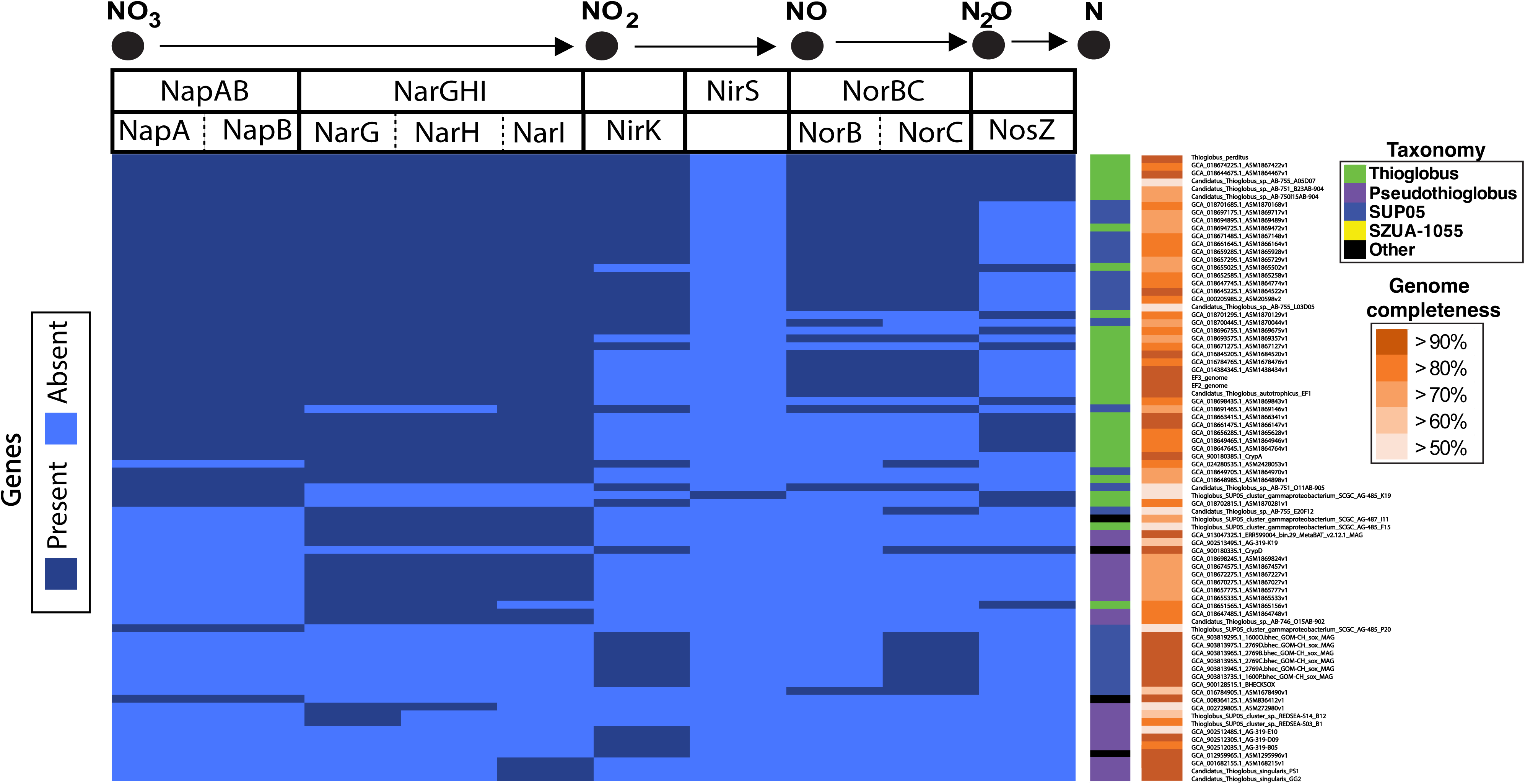
Dissimilatory denitrification pathway with only those genomes that encoded at least one step in this pathway (80/217) shown here. The presence or absence of genes involved in dissimilatory denitrification are represented by dark blue (present) and light blue (absent). Genome completeness is color-coded based on percent completeness. Further, the genomes are color-coded by taxonomic classification at the genus level. In the legend, black indicates the “other” category and shows clades represented by less than three genomes. The enzymes of each pathway are labeled at the top of the heatmap, along with a depiction of the pathway where black circles represent the products and/or intermediates. The enzymes with multiple subunits are divided by dashed lines. The enzymes are abbreviated as: nitrate reductase subunit A (NapA), nitrate reductase subunit B (NapB), respiratory nitrate reductase subunit G (NarG), respiratory nitrate reductase subunit H (NarH), respiratory nitrate reductase subunit I (Nar I), nitrite reductase subunit K (NirK), nitrite reductase subunit S (NirS), nitric oxide reductase subunit B (NorB), nitric oxide reductase subunit C, and nitrous oxide reductase (NosZ).

### Sulfur oxidation and reduction

Genome annotation revealed partial and complete pathways for dissimilatory sulfate energy metabolism, including both reduction and oxidation. In total, 91% (197 of 217) of the genomes encoded dissimilatory sulfate oxidation and reduction genes (Fig. 5A). Of the 197 genomes, 88 encoded all genes in complete sulfate oxidation and reduction, including sulfate adenylyltransferase (*sat*), adenylylsulfate reductase, subunit A (*aprA*), adenylylsulfate reductase, subunit B (*aprB*), dissimilatory sulfite reductase alpha subunit (*dsrA*), and dissimilatory sulfite reductase beta subunit (*dsrB*) (Fig. 5A). Specifically, these 88 genomes consisted of 70

**Figure 5.**
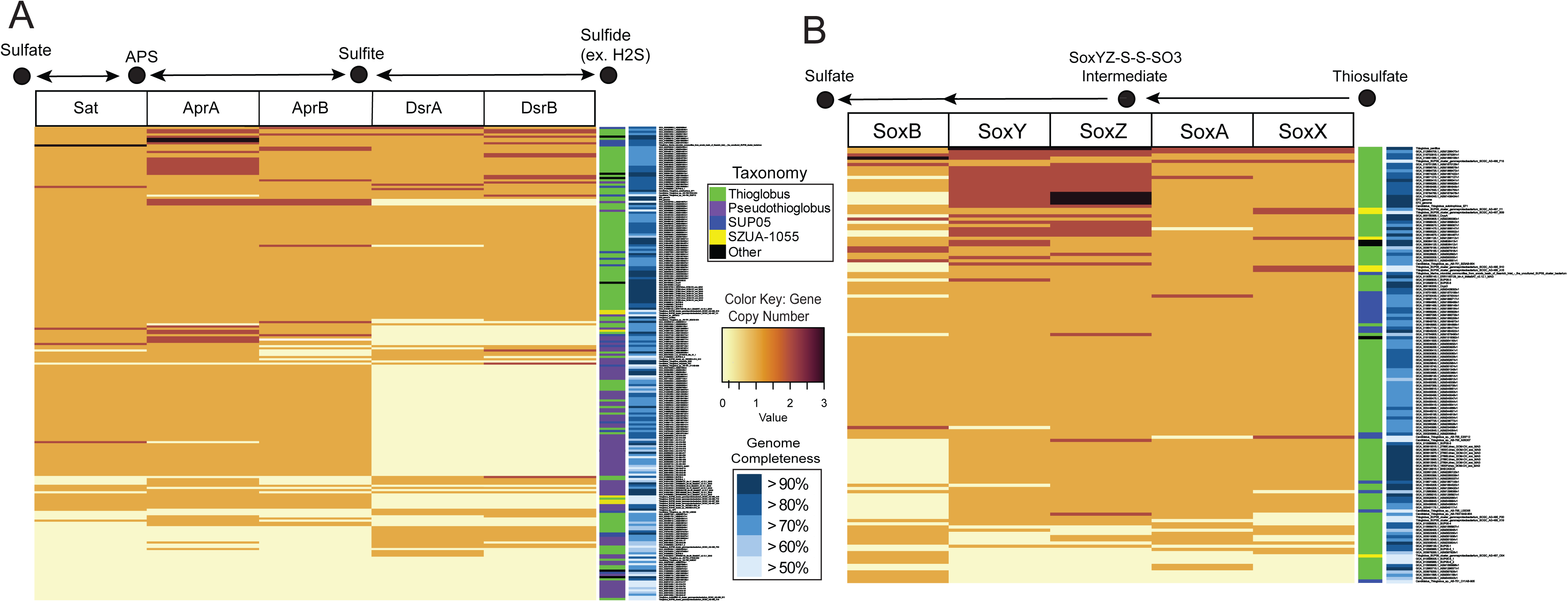
(A) Dissimilatory sulfate oxidation/reduction pathway and (B) sulfur oxidation via Sox enzyme complex with gene copy number. The number of gene copies per genome is indicated by the color key. Genomes are color-coded by taxonomic classification at the genus level. Black, indicating the “other” category represents clades represented by less than three genomes. Genome completeness is color-coded based on percent completeness. The Sox enzyme complex is abbreviated as: sulfate thiohydrolase (SoxB), sulfur oxidizing protein subunit Y (SoxY), sulfur oxidizing protein subunit Z (SoxZ), L-cysteine S-thiosulfotransferase subunit A (SoxA), L- cysteine S-thiosulfotransferase subunit X (SoxX). The enzymes for sulfate oxidation/reduction are abbreviated as: sulfate adenylyltransferase (Sat), adenylylsulfate reductase, subunit A (AprA), adenylylsulfate reductase, subunit B (AprB), dissimilatory sulfite reductase alpha subunit (DsrA), dissimilatory sulfite reductase beta subunit (DsrB). Black circles represent products and/or intermediates. Double arrows indicate that the steps in the pathway can proceed in either direction.

*Thioglobus*, 9 SUP05, 3 *Pseudothioglobus*, 3 *Thiodubiliella*, 2 SZUA-1055, and 1 JAAOIFI01. Additionally, 129 of the 197 genomes encoded an incomplete pathway, including 76 *Pseudothioglobus*, 37 *Thioglobus*, 10 SUP05, 4 SZUA-1055, and 2 DUCF01 (Fig. 5A).

Genes involved in the Sox multi-enzyme complex (SoxABXYZ) are representative metabolic markers for inorganic sulfur oxidation via the oxidation of thiosulfate to sulfate (Spring, 2014; Baltar et al., 2023). Gene annotations revealed that 63% (136 of 217) of the *Thioglobaceae* genomes encoded all essential *sox* genes for sulfur oxidation, including members of the genera SUP05, *Thioglobus*, SZUA-1055, *Thiodubiliella*, and JAAOIFI01 (Fig. 5B). Of the analyzed genomes, 73 encoded all *sox* genes required for sulfur oxidation, while 64 genomes coded for a subset of these genes (Fig. 5B). Specifically, 54 *Thioglobus*, 13 SUP05, 3 SZUA- 1055, two *Thiodubiliella*, and one JAAOIFI01 encoded the full Sox multi-enzyme complex (Fig. 5B). Given that sulfate thiohydrolase (*soxB*) is commonly used as a metabolic marker for sulfur oxidation, as it encodes for the final component of oxidizing thiosulfate to sulfate (Spring 2014; Baltar et al., 2023), it is notable that 93 genomes encoded the *soxB* gene.

## Discussion

### Dissolved Oxygen, Microbial Diversity and Abundance

As global ocean temperatures rise, DO levels are expected to decline, fueling the expansion of global OMZs (Keeling et al., 2010; Breitburg et al., 2018), with a growing understanding of how low DO impacts microbial community dynamics (Wright et al., 2012). Here, we investigated changes in diversity and community composition along a DO gradient in the nBUS, and identified DO as a primary driver shaping microbial community diversity and structure. For example, community diversity was significantly lower in suboxic samples as compared to oxic samples, which can largely be attributed to the low evenness in suboxic samples. Our findings are consistent with reports in which microbial diversity metrics were determined in OMZs. Specifically, community diversity in OMZs decreased within low DO environments (Bryant et al., 2012; Beman & Carolan, 2013). In our samples, community richness was highest in dysoxic samples, and significantly lower in both oxic and suboxic samples. This observation aligns with Beman and Carolan (2013), who reported a unimodal pattern between DO and bacterial richness, with a peak in richness at the edges of the OMZ and decreasing richness in the interior. This is also consistent with an OMZ off British Columbia, where a large decline in richness was observed when transitioning into a hypoxic environment (Zaikova et al., 2010), underscoring the inverse relationship between microbial richness and DO (Spietz et al., 2015).

The significant decrease in community evenness in suboxic samples compared to both oxic and dysoxic samples, suggested that DO regulates, in part, the abundance of particular microbes. This is supported by previous reports that revealed the dominance of distinct microbial groups in low DO samples (Beman & Carolan, 2013; Spietz et al., 2015; Aldunate et al., 2018), which would influence community evenness. Similarly, we found that the abundance of *Thioglobaceae* (and *Nitrosopumilaceae*) increased significantly as DO decreased. In fact, *Thioglobaceae* comprised up to 40% of the microbial community in the low DO samples. Our observations are consistent with other analyses of *Thioglobaceae*, which is reported as being a dominant member of the microbial community in several OMZs (Lavik et al., 2009; Walsh et al., 2009; Glaubitz et al., 2013; Callbeck et al., 2018; Morris & Spietz 2022), including the nBUS (Lavik et al., 2009; Vuillemin et al., 2022 ; Dangl et al., 2025)). Additionally, *Thioglobaceae* have been shown to be abundant at oxic-anoxic interfaces in OMZs by Morris and Spietz (2022). Finally, recent evidence suggested that some *Thioglobaceae* are optimized to grow at low DO concentrations (Mattes et al., 2021).

### Genomic Analysis of the Metabolic Function of Thioglobaceae

#### Carbon

Only SUP05 and *Thioglobus* encoded a complete carbon fixation pathway through the canonical CB cycle. The canonical CB cycle and the non-canonical CB cycle (NCB) are set apart by FBP and SBP hydrolysis activity, which is mediated by the bifunctional enzyme F/SBpase (Hudson et al., 2024). The canonical CB cycle includes F/SBpase, which was observed in less than 15% of the genomes. Most of the remaining genomes lacked F/SBpase, suggesting that the majority of *Thioglobaceae* carry out carbon fixation via a variant of the CB cycle, as all other components of the CB cycle were present. Possible variants for sulfur oxidizing chemoautotrophs is the replacement of F/SBpase with PPi-PFKs, which operate in reverse (Kleiner et al., 2012).

Notably, our analysis revealed that at least one representative of each genera within the *Thioglobaceae* coded for complete carbon fixation through either the CB or NCB pathway. Recently, *Ca. T. autotrophicus* strains EF2 and EF3 isolates were shown to fix inorganic carbon with the NCB pathway (Morris & Mino, 2024), that was originally reported in strain EF1 (Shah & Morris, 2015). This finding is significant given that the SUP05 clade (as it is referred to in the literature) has been reported to use the energy obtained from reducing inorganic sulfur to fuel autotrophic carbon fixation via RuBisCO (Walsh et al., 2009; Spietz et al., 2019; Morris & Spietz, 2022), which was identified in 78% of the genomes analyzed, including strains EF2 and EF3. This was verified through experiments with *Ca. T. autotrophicus* (strain EF1) where Shah and Morris (2015), showed that it uses energy from sulfur oxidation to fix inorganic carbon with RuBisCo.

The genomes of strains EF1, EF2, and EF3 derive from cultured representatives and are complete. All three have NCB pathways and it has also been demonstrated that strain EF1 requires inorganic carbon for growth. Overall, the ability to fix carbon either via the CB or NCB pathway was encoded in a large number of the analyzed genomes. These genomes spanned taxonomic groups, and represented microbes sampled from the global ocean, suggesting that *Thioglobaceae* as a whole are important contributors to the marine carbon cycle through carbon fixation.

Analysis of carbon utilization by *Thioglobaceae* also revealed that key components of the TCA cycle, such as 2-oxoglutarate dehydrogenase, were lacking in most genomes. This suggested that *Thioglobaceae* may utilize a “horseshoe TCA cycle,” an alternative TCA pathway that directs organic carbon into central carbon metabolism to conserve organic carbon for biosynthesis (Wood et al., 2004; Morris & Spietz, 2022). This cycle differs from the canonical TCA cycle, which is primarily for amino acid biosynthesis, not energy acquisition (Wood et al., 2004; Morris & Spietz, 2022).

#### Aerobic and Anaerobic Respiration

Low and high affinity oxygen enzymes, cytochrome c oxidase subunits I-III (CoxABC) and cbb3-type cytochrome c oxidase subunits I-II, were identified in nearly half of the analyzed genomes, indicating the capacity for aerobic respiration. The presence of both low and high affinity enzymes in these genomes suggests a potential adaptive mechanism for varying oxygen conditions.

Over a third of the genomes analyzed encoded at least one step in dissimilatory denitrification, which suggested anaerobic respiration. Of these, only six *Thioglobus* genomes encoded complete denitrification. Further, several genomes lacked genes coding for NosZ, the enzyme that mediates the last denitrification step. This denitrification pathway was truncated with nitric oxide reductase (NorBC), which converts NO to N₂O. Notably, 13 of the 14 genomes encoding incomplete denitrification belong to the genus SUP05, several of which were sampled from the Saanich Inlet (Lin et al., 2021). For example, *Ca. T. autotrophicus* (strain EF1) has been shown to evolve N₂O under anaerobic growth conditions (Shah et al., 2017). The nBUS has been reported to have elevated sea-air fluxes of N₂O and accounts for up to 13% of the total coastal upwelling N₂O source (Arévalo-Martínez et al., 2019). Taken together, the high abundance of SUP05 in OMZs, along with culture experiments involving *Ca. T. autotrophicus*, in which N₂O was produced from incomplete denitrification, suggested that this group could play an important role in greenhouse gas production in the marine environment. Additionally, new data presented herein showed that globally distributed *Thioglobaceae* lack the enzyme required to convert N₂O to N₂. Importantly, incomplete denitrification, such as that encoded by *Thioglobaceae*, was shown to increase N₂O cycling rates by an order of magnitude over modeled estimates in the suboxic Eastern Tropical North Pacific (Babbin et al., 2015).

The BUS is a known source of N₂O (Nevison et al., 2004; Frame et al., 2014, Arévalo-Martínez et al., 2019; Dangl et al., 2025). In the nBUS Frame et al. (2014) reported fluxes as high as 500 mg N/m2/yr. They reported the isotopic composition of N₂O at the maxima was largely the result of denitrification and/or nitrifier denitrification. Dangl et al. (2025) sampled four stations in the nBUS during the austral winter and reported the N₂O isotopic composition in low oxygen samples was predominantly the result of denitrification. Further, they assembled one *Thioglobaceae* MAG that encoded an incomplete denitrification pathway, as it lacked *nosZ,* and suggested that it may be an important denitrifier potentially involved in N₂O production in the nBUS. When these observations from the environment, including our work presented herein, are coupled with experiments with *Ca. T. autotrophicus* showing N₂O production (Shah et al., 2017), a better understanding of *Thioglobaceae* N₂O production emerges and suggests that this clade may be important contributors to N₂O emissions in the nBUS. Further, the inclusion of *Thioglobaceae* genomes from the global ocean in our study expands our findings to other OMZs.

#### Sulfur oxidation and reduction

We identified several SUP05 and *Thioglobus* genomes containing key enzymes in known sulfur oxidation pathways that have the potential to oxidize a range of reduced sulfur compounds. Specifically, *Thioglobus,* SUP05, and a small number of other *Thioglobaceae* genomes encoded essential *sox* genes, which would enable complete sulfur oxidation. The presence of the Sox multi-enzyme complex would allow for energy acquisition from the oxidation of thiosulfate, elemental sulfur, sulfide, and sulfate (Baltar et al., 2023). SUP05 displays a high affinity for sulfide (Crowe et al., 2018) and are recognized as key contributors to sulfide oxidation in OMZs (Long et al., 2021). Further, 40% of the genomes, predominantly classified as *Thioglobus,* encoded the complete pathways for both dissimilatory sulfate reduction and oxidation, including the key genes dissimilatory sulfite reductase (d*srAB*), adenylylsulfate reductase (*aprAB*), and sulfate adenylyltransferase (*sat*). Further, *Thioglobus* and SUP05 in particular, have the genetic potential to use the energy gained from reducing inorganic sulfur to drive carbon fixation. Given that SUP05 and *Thioglobus* are among the most abundant microbes in OMZs, their role in sulfur oxidation may be ecologically consequential. For instance, toxic hydrogen sulfide (H_2_S) is reported to accumulate in the nBUS (Callbeck et al., 2018). High abundances of SUP05 in the nBUS are partly credited with the detoxification of these sulfidic waters (Lavik et al., 2009), which have contributed to massive fish kills off the African coast (Copenhagen, 1953; Hamukuaya *et al.*, 1998).

Collectively, our analyses revealed that DO influenced microbial community diversity and structure, with significant increases in the abundance of *Thioglobaceae,* as DO concentrations declined in the nBUS. This data also revealed the dominance of *Thioglobaceae* in the nBUS along an oxygen gradient. Comparing iTag and genomic datasets suggested that the abundant *Thioglobaceae* were captured in the nBUS and global ocean genomes analyzed here.

Genomic analysis revealed that *Thioglobaceae*, sampled from the global ocean, have the genetic capacity to be important mediators of carbon, nitrogen, and sulfur cycling in the marine environment. For example, *Thioglobus* and SUP05 genomes encoded the necessary genes used to fix carbon via the canonical CB pathway, while other *Thioglobaceae* may do so by a NCB pathway. Further, the SUP05 genomes encoded low and high affinity enzymes for aerobic respiration, and also for denitrification. This suggested that *Thioglobaceae* are potentially metabolically active across an oxygen gradient, from oxic to suboxic, using different metabolic strategies for respiration. Importantly, when oxygen concentrations are low and *Thioglobaceae* increases in abundance, the lack of genetic machinary to convert N₂O to N₂ that was observed in several genomes suggested that this group could be an important contributor to N₂O flux from OMZs to the atmosphere. Finally, while *Thioglobus* and SUP05 genomes encode both SOX and sulfate oxidation and reduction pathways, this was also observed in other *Thioglobaceae* members, but with varying degrees of pathway completion. Thus, declining DO concentrations in the global ocean would be met with increasing abundances of a microbial group capable of fixing carbon dioxide and detoxifying the sulfidic water column, while also producing a potent greenhouse gas through incomplete denitrification.

## Materials and Methods

### Study Site and Data Collection

Seawater was collected from 11 coastal stations in the nBUS on the R/V Mirabilis in April 2017, during austral fall (Table 1, Fig. 1A). A total of 46 samples were collected at various depths (ranging from 2 to 972 m) across an oxygen gradient, which was classified into three categories: oxic (>2.0 ml/L), dysoxic (0.2–2.0 ml/L), and suboxic (0.0–0.2 ml/L) (Algeo and Li, 2020) (Table 1). At each station 5-10 L of seawater was collected at each depth. Seawater was pre-filtered using a 2.7 μm filter and concentrated on a 0.22 μm Millipore Sterivex filter.

**Table 1.**
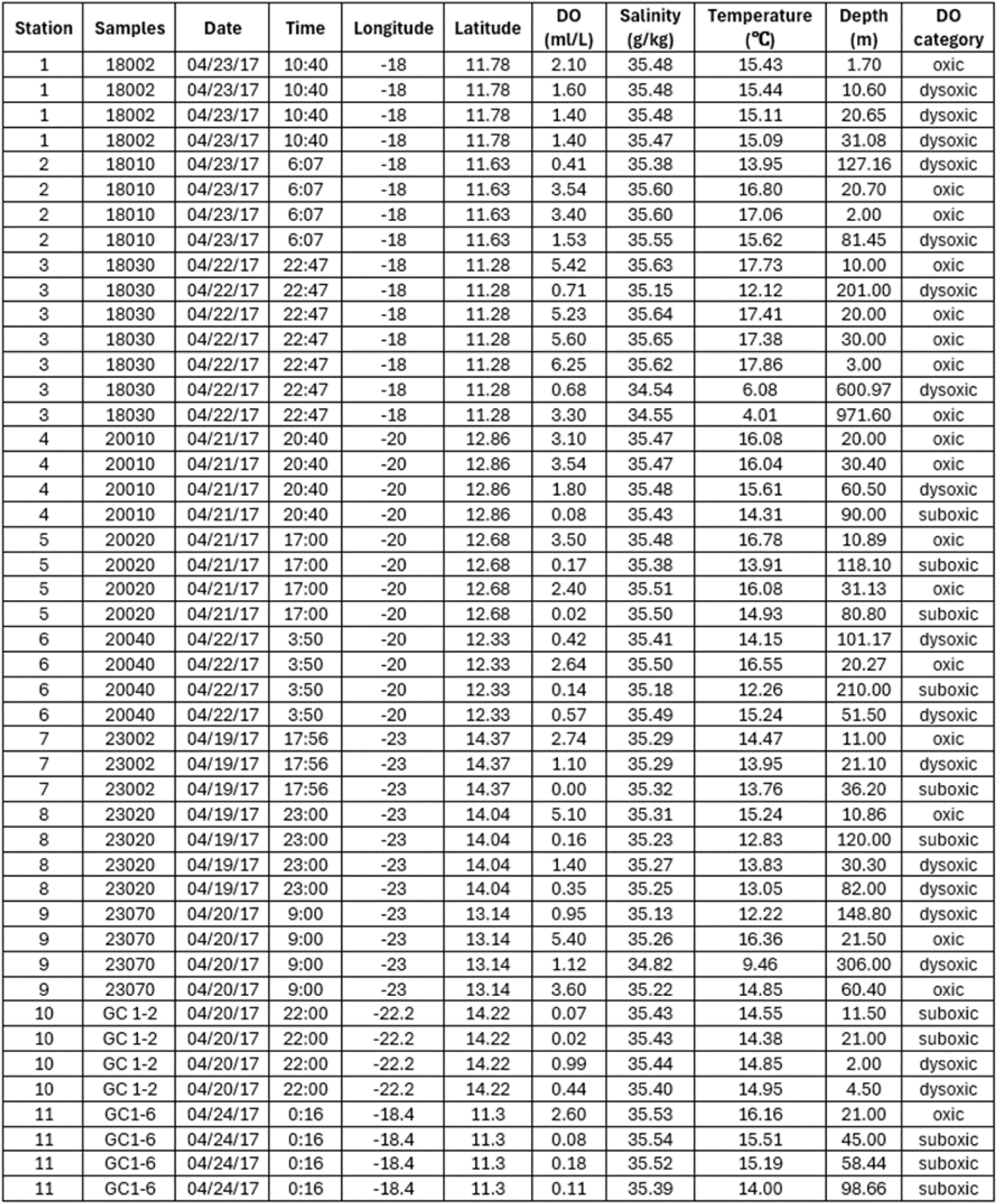
Metadata and environmental data for 11 stations sampled in the nBUS.

RNAlater was added to the Sterivex filter and it was stored at -80℃ until DNA extraction.

### DNA Extraction and 16S rRNA Gene Sequencing (iTag)

DNA was extracted from one half of the Sterivex filter, following the protocol described in Gillies et al. (2015). Briefly, DNA was extracted using a CTAB extraction buffer ((10% CTAB (hexadecyltrimethylammonium bromide), 1M NaCl, and 0.5 M phosphate buffer, pH 8) with 0.1M ammonium aluminum sulfate, and 25:24:1 phenol:chloroform:isoamyl alcohol) paired with bead beating via FastPrep-24 (MP Biomedicals). Genomic DNA was purified with a QIAGEN AllPrep DNA/RNA Kit (Valencia, CA, USA). The purified DNA was quantified using a Qubit2.0 Fluorometer (Life Technologies, Grand Island, NY, USA).

Microbial community structure was determined by sequencing the V4 region of 16S rRNA genes (hereinafter referred to as iTag) in all 46 samples. Specifically, 16S rRNA genes were amplified from 10ng of purified genomic DNA in duplicate using the modified archaeal and bacterial primers 515FB and 806RB (these primers target the V4 region) (Apprill et al., 2015; Parada et al., 2015) in accordance with the protocol described by Caporaso et al. (2011, 2012) and used by the Earth Microbiome Project (http://www.earthmicrobiome.org/emp-standard-protocols/16s/), with a slight modification: the annealing temperature was 60 °C. Amplicons were sequenced using an Illumina MiSeq in 250 × 250 bp mode.

### Bioinformatics, Taxonomy and Statistical Analyses of 16S rRNA Gene Sequences

Raw sequences were demultiplexed using QIIME2 (Caporaso *et al.*, 2010; ver. 2018.6). Demultiplexed reads were quality filtered, including chimera removal, and joined using DADA2 (Callahan *et al.*, 2016) using default parameters. The resulting amplicon sequence variant (ASV) (Callahan *et al.*, 2017) table was normalized using the Wrench package with the W2 estimator and defaults as recommended (Kumar et al., 2018). ASV taxonomy was assigned with the GTDB database (Parks et al., 2020); release 226) in QIIME2 using classify-sklearn. ASV data was normalized using multiple rarefactions, which were then used to determine Shannon diversity (Shannon and Weaver, 1949), richness (Chao1; Chao, 1984) and evenness (Pielou e; Pielou, 1966) using QIIME2. To test for significant differences in alpha diversity metrics and normalized ASV abundances between samples with different DO concentrations a Kruskal-Wallis rank sum test with the B-H correction was used.

The Wrench normalized ASV table was analyzed using non-metric multidimensional scaling (NMDS) in R with the metaMDS command in the Vegan package (Oksansen et al., 2019). Environmental variables and alpha diversity values were analyzed in relationship to NMDS axes with 999 permutations of the data The simper command (similarity percentage (SIMPER)) was used to evaluate the contribution of ASVs to Bray-Curtis dissimilarity, with oxic, dysoxic, and suboxic groupings. This was followed by a Dunn test, which is a nonparametric pairwise multiple comparisons statistic (Dinno, 2015), to test for significant differences in abundance based on DO concentrations. The effect of DO concentrations on microbial community composition was tested using the distance matrix with permutational multivariate analysis of variance (PERMANOVA) with the *adonis2* command. The ‘pairwiseAdonis’ package (Martinez Arbizu, 2020) was used to carry out multi-level pairwise comparisons with p-values adjusted with FDR. Beta-dispersion (“betadisper”, type=centroid, permutest with 999 permutations) was not significantly different between the treatments.

### Genome Similarity, Classification and Gene Annotation

The Integrated Microbial Genomes Database (IMG) and Genome Taxonomy Database (GTDB) were used to search for all SUP05, as this group has been historically named, and more generally, for all genomes in the *Thioglobaceae* family. This search resulted in 324 publicly available genomes identified in and downloaded from IMG or identified in GTDB and downloaded from the National Center for Biotechnology Information (NCBI). CheckM was used to identify genome completeness and examine percent genome contamination. In total, 217 genomes were identified that met the criteria of medium (>50% complete and <10% contamination) to high quality (>90% complete and <5% contamination). These genomes represent both cultured and uncultured microbes. Taxonomy was assigned to each genome using the Genome Taxonomy Toolkit (GTDB-Tk, ver. 2.3.0; with reference to GTDB R08-RS214) (46) with up to 120 bacterial single-copy marker proteins. Genomes were annotated using DRAM (ver. 1.1.1) (47) with the KEGG (48), UniRef90 (49), PFAM (50), dbCAN (51), and MEROPS (52) databases. Blastn was used to compare 16S rRNA gene iTag data from the nBUS dataset to those genes in the genomes analyzed here. Average nucleotide identity (ANI) was calculated using sourmash (Brown and Irber, 2016) with the following settings: sketch dna -p scaled=1000,k=21,k=31,k=51 and compare --containment --ani --ksize 31 to compare genetic relatedness among prokaryotic strains.

## Acknowledgments

We gratefully acknowledge the assistance of the instructors and students of the Regional Graduate Networks of Oceanography Discovery Camps, as well as the scientific staff and the crew on the R/V Mirabilis who made access to the Benguela Current Ecosystem possible. The samples were collected during the Mirabilis cruise in April 2017. The Discovery Camps of the Regional Graduate Network for Oceanography are funded by the Agouron Institute, the Simons Foundation, the Scientific Committee for Oceanographic Research, the Ministry of Fisheries and marine Resources through the National Marine Information and Research Center the University of Namibia, and ETH Zurich.

## Availability of Data and Materials

All 16S rRNA sequences are available through NCBI (PRJNA1251143), and on the Mason lab server http://mason.eoas.fsu.edu in the nBUS directory. Publicly available genomes were downloaded from IMG and NCBI.

